# Structure-based stabilization of SOSIP Env enhances recombinant ectodomain durability and yield

**DOI:** 10.1101/2022.10.25.513757

**Authors:** Daniel Wrapp, Zekun Mu, Bhishem Thakur, Katarzyna Janowska, Oluwatobi Ajayi, Maggie Barr, Robert Parks, Beatrice H. Hahn, Priyamvada Acharya, Kevin O. Saunders, Barton F. Haynes

## Abstract

The envelope glycoprotein (Env) is the main focus of HIV-1 vaccine development due to its critical role in viral entry. Despite advances in protein engineering, many Env proteins remain recalcitrant to recombinant expression due to their inherent metastability, making biochemical and immunological experiments impractical or impossible. Here we report a novel proline-stabilization strategy to facilitate the production of prefusion Env trimers. This approach, termed “2P”, works synergistically with previously described SOSIP mutations and dramatically increases the yield of recombinantly expressed Env ectodomains without altering the antigenic or conformational properties of native Env. We determined that the 2P mutations function by enhancing the durability of the prefusion conformation and that this stabilization strategy is broadly applicable to evolutionarily and antigenically diverse Envs. These findings provide a new Env stabilization platform to facilitate biochemical research and expand the number of Env variants that can be developed as future HIV-1 vaccine candidates.

## INTRODUCTION

HIV-1 infection causes severe CD4^+^ T cell immunodeficiency that is accompanied by fever, weight loss and opportunistic infections(1). Recent estimates place the number of new infections at approximately 1.5 million per year(2). Despite decades of rigorous research and life-saving advances in both pre- and post-exposure anti-retroviral drug treatments(3, 4), the development of an effective HIV-1 vaccine remains an unmet public health goal(5).

HIV-1 is a lentivirus that uses a class I viral fusion glycoprotein, called envelope (Env), to gain entry into host cells to begin the process of integration and replication(6). Like other class I viral fusion proteins, Env is proteolytically processed into two subunits, gp120 and gp41(7). These subunits remain associated and oligomerize with other protomers to form the functional prefusion conformation of Env, which is composed of a trimer of gp120/gp41 heterodimers. The N-terminal gp120 subunit is responsible for mediating binding to both the CD4 receptor and CCR5/CXCR4 coreceptor(8-10), while the gp41 subunit contains the hydrophobic fusion peptide and the helical heptad repeats that drive membrane fusion(11).

Because Env is the sole target for neutralizing antibodies, it is currently the main focus of vaccine research. However, several characteristics of Env make it a notoriously complex and problematic immunogen. These include a dense glycan shield composed of *N*-linked glycans that cover the ectodomain and hamper the elicitation of neutralizing antibodies(12). Furthermore, high levels of viral replication, constant immune pressure(13) and the infidelity of the HIV-1 reverse transcriptase have resulted in extreme Env sequence diversity among viral strains(14). Another problem is the conformational heterogeneity that Env is capable of displaying. The prefusion, closed conformation that Env adopts prior to host-cell receptor engagement is immunologically ideal for the elicitation of broadly neutralizing antibodies (bnAbs)(15). In this conformation, Env presents bnAb epitopes while simultaneously concealing many of the non-neutralizing epitopes which elicit an unproductive immune response(16). However, Env has evolved to be metastable in this prefusion conformation, such that it is primed to rapidly and irreversibly transition to the postfusion conformation(17). This inherent metastability makes it difficult to recombinantly express the Env ectodomain in the antigenically desirable prefusion conformation. In an effort to stabilize the prefusion conformation of the Env ectodomain without altering its antigenicity, Sanders et al. developed the SOSIP mutations, composed of an engineered disulfide bond (SOS) that links the gp120 and gp41 subunits and an I559P (IP) substitution that disfavors the formation of the elongated postfusion helices that make up the six-helix bundle(18, 19). Since their initial description, the SOSIP mutations have undergone iterative improvement to enhance the thermostability and the antigenic characteristics of the prefusion trimer(15, 20, 21). Alternative and complimentary protein engineering approaches have also been described as a means of yielding improved Env immunogens(22-26). However, despite these significant advances in immunogen engineering, many Env variants remain recalcitrant to recombinant *in vitro* expression.

Here we describe a novel set of proline substitutions that increase the yield of recombinantly expressed prefusion Env. These substitutions, referred to as “2P”, function synergistically with previously reported SOSIP mutations and do not alter the antigenicity or the overall structure of the stabilized Env trimer. We go on to show that the mechanism by which these mutations increase yield is through enhancing the durability of the prefusion conformation, rather than through boosting expression levels. Moreover, we show that the 2P mutations can effectively be applied to a broad range of antigenically and evolutionarily diverse Envs. By facilitating the expression and purification of soluble, near-native Envs, the 2P mutations should serve as a useful tool for investigating medical countermeasures against HIV-1.

## RESULTS

In an effort to enhance the yield of recombinantly expressed, stabilized Env trimers, we designed a series of single proline substitutions throughout the truncated gp41 subunit. Proline substitutions were specifically evaluated due to their propensity to act as “helix breakers” since their cyclized side chain prevents conventional amide backbone hydrogen bonding(27). Based on structural analysis of both the prefusion and postfusion conformations of gp41, nine positions were selected to disfavor the formation of the elongated alpha helices which are characteristic of class I fusion proteins in the postfusion state (**Fig. 1A**). These substitutions were then made in the CH848 10.17DT Env, a clade C Env that has been modified to engage the unmutated common ancestor (UCA) from the DH270 V3 glycan bnAb lineage(28, 29). This Env was expressed using a chimeric gp41 subunit from BG505 and was stabilized with both SOSIP.664 and DS mutations (CH848 10.17DT DS-SOSIP)(18, 24, 30). Purification of these constructs after transient transfection revealed that several of the individual proline substitutions had a dramatic effect on Env yield (**Fig. 1B**). Envs with proline substitutions at positions 536, 545, 568 or 569 (HXB2 numbering) in particular showed a marked increase in the yield of prefusion trimer relative to the unmodified SOSIP construct. Therefore these four mutations were selected for further analysis. Interestingly, although they were not incorporated into the final construct, proline substitutions at positions 545 and 569 were previously tested by Sanders et al. during the initial design of the SOSIP mutations. However, these mutations were deemed less effective than I559P in the context of the BG505 Env and were never evaluated in combination with the I559P substitution.

**Figure 1:**
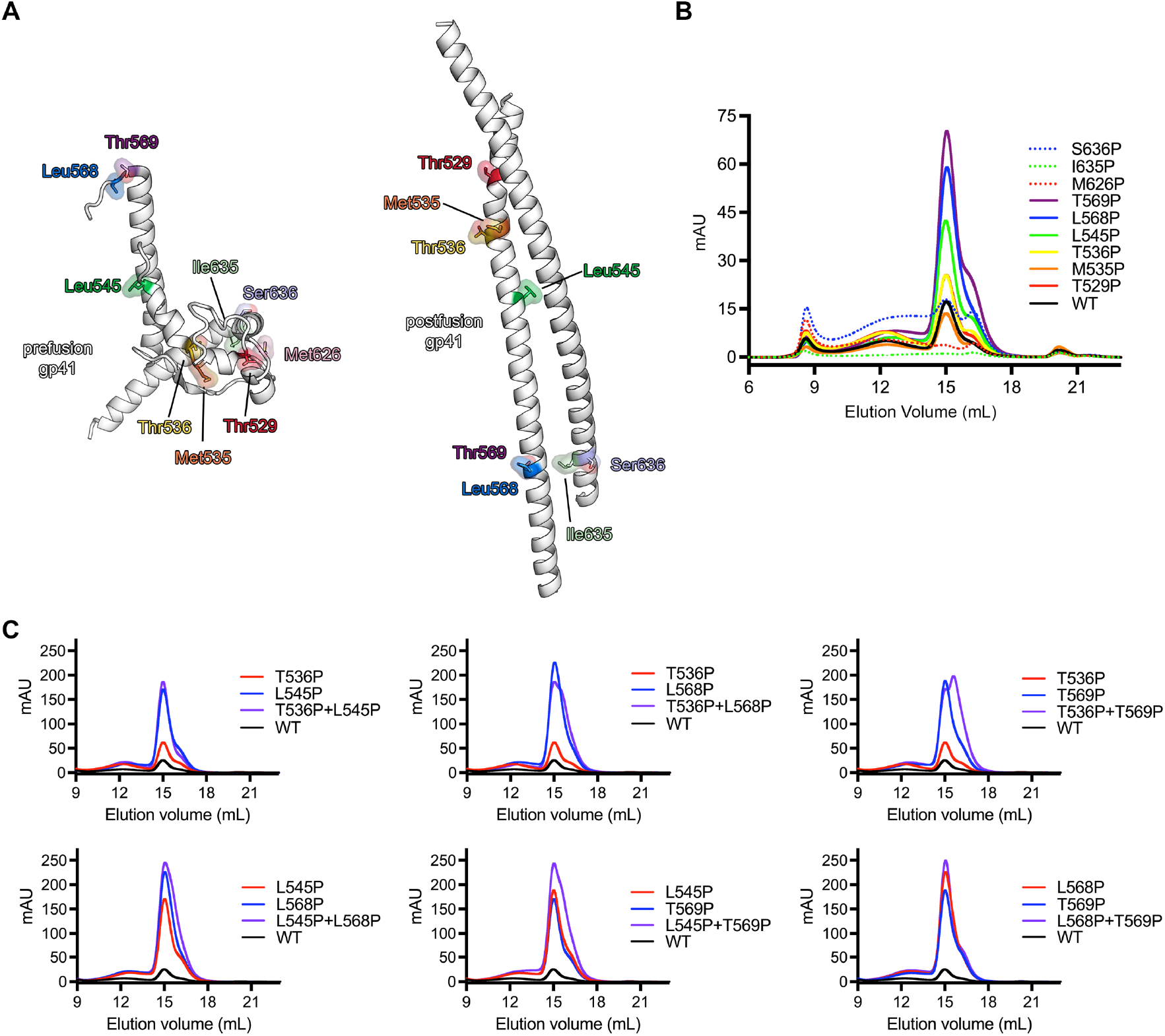
Design and evaluation of novel proline-stabilization mutations in gp41. (**A**, *left*) A single monomer of gp41 in the prefusion conformation (PDB ID: 6VZI) is shown as ribbons, colored white. Residues that were targeted for proline stabilization are shown as colored sticks surrounded by a transparent molecular surface. (**A**, right) A single monomer of gp41 in the postfusion conformation (homology model based on PDB ID: 7AEJ) is shown, colored according to the prefusion monomer. (**B**) Size-exclusion chromatograms from a Superose 6 Increase column for each individual mutant after affinity chromatography are overlaid. (**C**) Size-exclusion chromatograms from a Superose 6 Increase column are shown for individual proline mutants, colored either blue or red. The combination of the two mutations is colored purple and the unmutated CH848 10.17DT DS-SOSIP is colored black and labeled “WT”.

Individual proline mutations at these four positions were combined to form double-mutants to determine whether these substitutions might be capable of functioning synergistically (**Fig. 1C**). Modest improvements in yield over the single substitutions were observed after combining substitutions 536+545 or 568+569, while other combinations resulted in either decreased yield (536+568) or the appearance of a prominent, contaminating low-molecular-weight species which was detectable by size-exclusion chromatography (536+569, 545+568, 545+569). Similarly, combining proline substitutions at all four positions did not result in enhanced yield as compared to the double mutant at positions 568 and 569 (**S.Fig. 1**), which resulted in an ∼8-fold increase in prefusion trimer relative to CH848 10.17DT DS-SOSIP. Therefore, the double mutant at positions 568 and 569 was selected for subsequent characterization and these mutations were termed “2P” to reflect their similarity to the two consecutive proline mutations which have been shown to stabilize betacoronavirus spike proteins in the prefusion conformation(31).

Intriguingly, the 2P mutations at positions 568-569 in gp41 are also positioned similarly to the 2P mutations in the S2 subunit of the coronavirus spike (**S.Fig. 2A**). Both sets of mutations cap the N-terminus of the HIV-1 gp41 or CoV S2 central helix, which ultimately polymerizes into an elongated helix during the transition from prefusion to postfusion. Generally the residues upstream of this helical capping position in gp41 cannot be clearly resolved during structural characterization, presumably due to a high degree of conformational flexibility. However, the recently reported structure of Env in the occluded-open conformation shows these amino acids forming a 5-residue helical extension of the central helix (**S.Fig 2B**), suggesting that the 2P mutations may be stabilizing the prefusion conformation of gp41 by disfavoring the sampling of this early intermediate(32).

**Figure 2:**
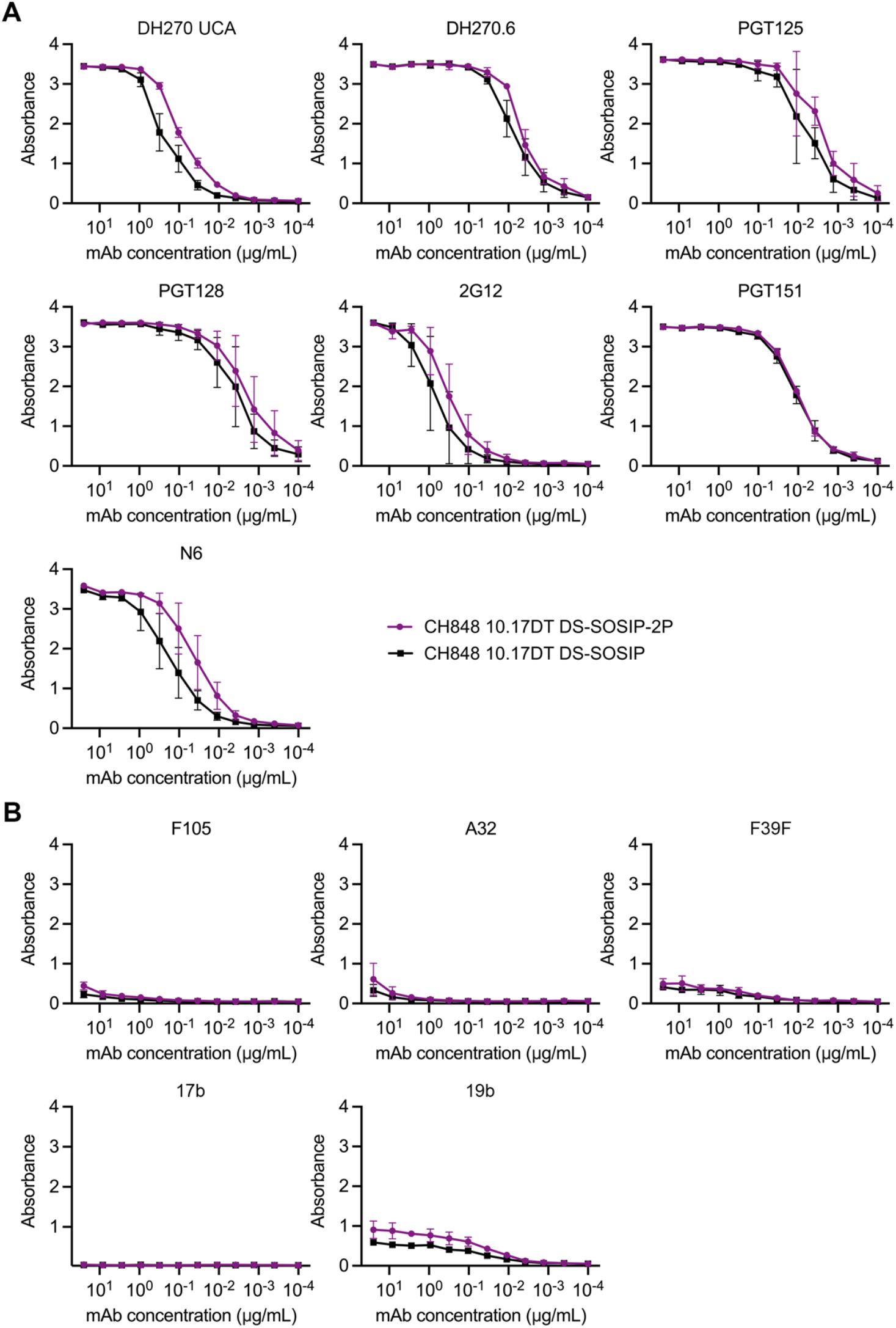
2P mutations do not alter the antigenic landscape of CH848 10.17DT DS-SOSIP. ELISA binding curves for Env-directed monoclonal antibodies against CH848 10.17DT DS-SOSIP and CH848 10.17DT DS-SOSIP-2P are shown. CH848 10.17DT DS-SOSIP is shown in black and CH848 10.17DT DS-SOSIP-2P is shown in purple. Data points represent the average of three replicates and the standard deviations are plotted as error bars. The y-axes show absorbance measured at 450 nm. Neutralizing antibody curves are shown in (**A**) and non-neutralizing antibody curves are shown in (**B**).

We next generated an “SOS 2P” Env by reverting the I559P mutation to investigate whether the novel 2P mutations could increase the yield of prefusion Env in the absence of the stabilizing effect of the IP mutation. The SOS 2P construct yielded only slightly more prefusion trimer than the SOSIP Env, while the dramatic boost in yield that was observed previously could only be recapitulated when the IP and 2P mutations were combined (**S. Fig. 3**). This synergistic stabilizing effect is reminiscent of the phenomenon that has been reported for the HexaPro mutations in the context of the SARS-CoV-2 spike protein, which further enhanced the effects of the initial S-2P mutations by engineering four additional proline substitutions throughout the S2 subunit(33).

**Figure 3:**
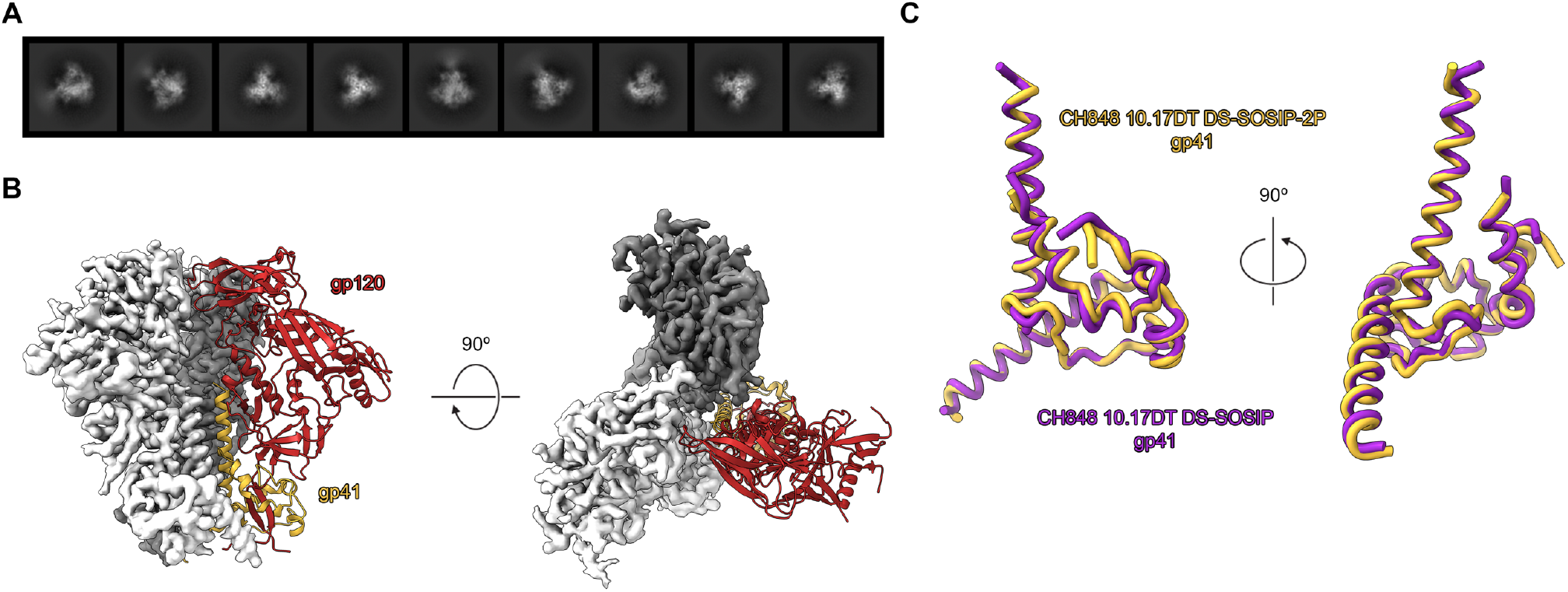
The cryo-EM structure of CH848 10.17DT DS-SOSIP-2P. (**A**) 2D class averages of the CH848 10.17DT DS-SOSIP-2P Env ectodomain, calculated in cryoSPARC v3, are shown. (**B**) The 3.73-Å-resolution reconstruction is shown from side (*left*) and top (*right*) views. Two protomers are displayed as the cryo-EM map in either white or dark gray and the third protomer is shown as a ribbon diagram of the corresponding model, with gp120 colored red and gp41 colored yellow. (**C**) A single monomer of gp41 from CH848 10.17DT DS-SOSIP-2P is shown in yellow, aligned to a monomer of gp41 from a previously determined cryo-EM structure of CH848 10.17DT DS-SOSIP (PDB ID: 6UM7), colored purple.

To determine whether the 2P mutations altered the antigenic landscape of the Env ectodomain, we performed an ELISA to compare the binding profiles of CH848 10.17DT DS-SOSIP and CH848 10.17DT DS-SOSIP-2P against a panel of well-characterized monoclonal antibodies (mAbs). Both neutralizing and non-neutralizing mAbs spanning multiple epitopes were included to comprehensively evaluate what effect the 2P mutations might have on Env folding (**Fig. 2**). Overall, the binding characteristics exhibited by the two Envs were very similar, although the 2P-stabilized construct appeared to bind slightly better to some neutralizing mAbs directed against the V3 glycan epitope (DH270 UCA) and the CD4-binding site (N6) (**Fig 2A**). To further validate that the newly introduced 2P mutations did not disrupt the overall folding of the gp41 subunit, we determined the cryo-EM structure of the CH848 10.17DT DS-SOSIP-2P Env to a resolution of 3.7 Å (**Fig. 3A-B, S.Fig. 4, S.Fig. 5, S.Table 1**). Our model spanned residues 32−664 and we were able to build *N*-linked glycans at 15 of the 25 putative sequons. There were several small, flexible loops in the gp120 subunit (59−66, 458−459, 400−411) that could not be confidently modeled, but overall the CH848 10.17DT DS-SOSIP-2P Env was virtually indistinguishable from the structures of previously reported CH848 10.17DT Envs which lack 2P stabilization(28). Like most other previously reported structures of gp41s in the prefusion conformation, the flexibility at the N-terminus of the central helix precluded us from observing both of our newly introduced proline substitutions, and only Pro569 could be confidently modeled into our reconstruction. However, the overall RMSD between our 2P-stabilized Env and a previously determined structure of CH848 10.17DT DS-SOSIP was only 0.757 Å over 1,253 Cα atoms (**Fig. 3B**), confirming that the 2P mutations are capable of enhancing the yield of recombinantly expressed Env ectodomain without altering the conformation of the SOSIP trimer.

**Figure 4:**
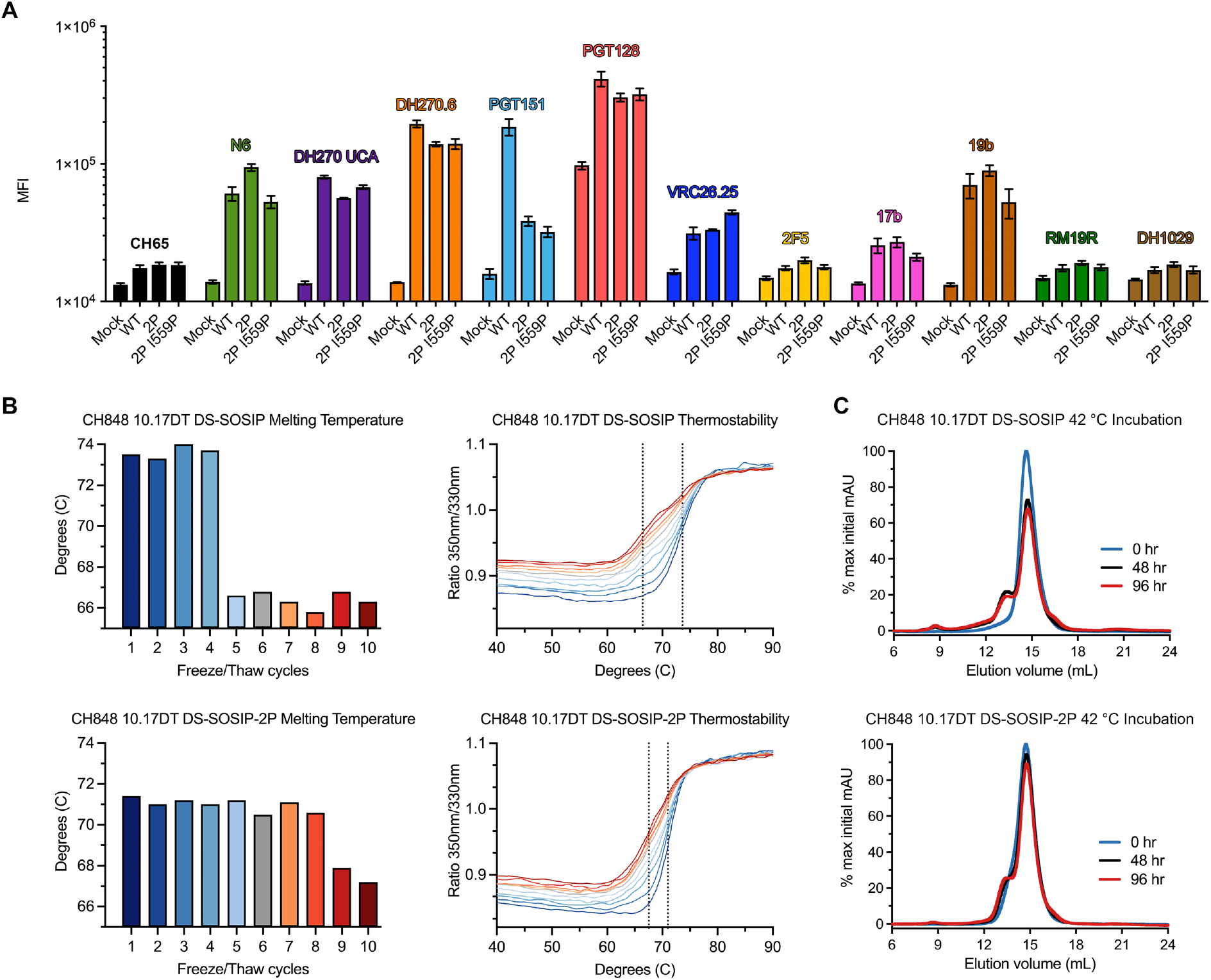
Investigating the underlying mechanism of enhanced protein yield upon proline-stabilization. (**A**) Four cell populations were stained with a panel of Env-directed monoclonal antibodies. The average of three independent experiments is plotted ± standard error of mean. Each antibody has been assigned a color and is labeled above the corresponding bars. “Mock” = untransfected, “WT” = CH848 10.17DT gp160, “2P” = CH848 10.17DT 2P gp160, “2P I559P” = CH848 10.17DT 2P I559P gp160. (**B**, *top left*) CH848 10.17DT DS-SOSIP Tm is plotted for each round of freeze/thaw. (**B**, *top right*) DSF melting curves used to calculate CH848 10.17DT DS-SOSIP Tm are plotted, colored according to the bar graph on the *left*. The dotted line at 73.6 °C represents the average Tm of rounds 1-4 and the dotted line at 66.4 °C represents the average Tm for rounds 5-10. (**B**, *bottom left*) CH848 10.17DT DS-SOSIP-2P Tm is plotted for each round of freeze/thaw. (**B**, *bottom right*) DSF melting curves used to calculate CH848 10.17DT DS-SOSIP-2P Tm are plotted, colored according to the bar graph on the *left*. The dotted line at 71.0 °C represents the average Tm of rounds 1-8 and the dotted line at 67.5 °C represents the average Tm for rounds 9-10. (**C**, *top*) Size-exclusion chromatograms for three aliquots of CH848 10.17DT DS-SOSIP, incubated at 42 °C for 0, 48 or 96 hours, are overlaid. The 0 hour sample is colored blue, the 48 hour sample is colored black and the 96 hour sample is colored red. Data are plotted as a percentage of the maximum mAU value from the 0 hour sample. (**C**, *bottom*) Size-exclusion chromatograms for three aliquots of CH848 10.17DT DS-SOSIP-2P, incubated at 42 °C for 0, 48 or 96 hours, are overlaid. The 0 hour sample is colored blue, the 48 hour sample is colored black and the 96 hour sample is colored red. Data are plotted as a percentage of the maximum mAU value from the 0 hour sample.

**Figure 5:**
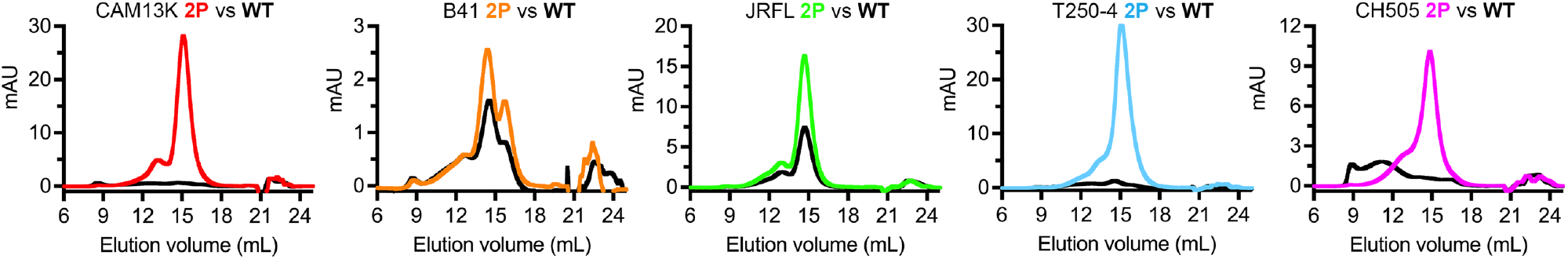
Applying 2P mutations to evolutionarily diverse Envs. Size-exclusion chromatograms are shown for each construct after PGT145 affinity chromatography. Curves for the 2P-stabilized construct are colored and the curves for the corresponding non-2P-stabilized constructs (“WT”) are overlaid in black. Chromatograms were generated using a Superose 6 Increase column. Full details about these Env constructs can be found in **S. Table 2**.

We hypothesized that this increase in the yield of prefusion trimer could either be due to an increase in Env expression or due to enhanced durability of the prefusion conformation upon proline-stabilization. To distinguish between these two possibilities, we began by investigating what impact proline-stabilization might have on cell-surface expression of full-length Env. FreeStyle 293-F cells were transiently transfected with CH848 10.17DT gp160s containing either no proline mutations, 2P mutations, or 2P mutations in combination with the I559P mutation. After 48 hours, the relative expression level of each membrane-anchored Env was assessed by flow cytometry after staining with a panel of mAbs. Despite the dramatic differences in yield that could be observed when evaluating the recombinant expression of soluble ectodomains, no significant differences in surface expression levels of membrane-anchored Envs could be detected among the three gp160 constructs that were tested (**Fig. 4A, S. Fig. 6**). The only clear difference between the three gp160s was a decrease in PGT151 binding in the 2P and the 2P+I559P constructs relative to the unstabilized gp160. However, this trend was not conserved for the rest of the antibody panel, nor was there a detectable difference between the 2P-stabilized and non-2P-stabilized Envs when testing PGT151 binding to soluble ectodomains (**Fig 2A**).

Based on the cell-surface expression results, we next evaluated the durability of recombinantly expressed, proline-stabilized Env ectodomains. Aliquots of CH848 10.17DT DS-SOSIP and CH848 10.17DT DS-SOSIP-2P were subjected to ten rounds of rapid freezing and thawing. Thermostability measurements of both Envs were collected by differential scanning fluorimetry (DSF) during each round (**Fig. 4B**). The melting temperature (Tm) of CH848 10.17DT DS-SOSIP began at 73.6 °C, a value that was maintained until after four freeze/thaw cycles, when it abruptly dropped to 66.4 °C, where it remained throughout the rest of the experiment. The Tm of CH848 10.17DT DS-SOSIP-2P was initially measured to be 71.0 °C, a value that was maintained until after eight freeze/thaw cycles, when the Tm dropped to 67.5 ° C. To further investigate this phenomenon, we performed a forced degradation assay in which aliquots of CH848 10.17DT DS-SOSIP and CH848 10.17DT DS-SOSIP-2P were incubated at 42 °C for either 48 or 96 hours. The integrity of the incubated trimers was then evaluated by size-exclusion chromatography (**Fig. 4C**). After 48 hours at 42 °C, we observed the appearance of a high-molecular weight aggregate peak in the non-2P-stabilized sample. The emergence of this peak corresponded with a 26.8% decrease in the amount of properly folded trimer, as measured by quantifying the area under the SEC curve. A similar, albeit less prominent, peak was not observed in the 2P-stabilized sample until after 96 hours of incubation at 42 °C, and the amount of properly folded trimer decreased by only 4.3%. These data, in conjunction with our evaluation of cell-surface expression levels, suggest that proline-stabilization of the gp41 subunit has little impact on the level of protein expression, and that observed increases in the yield of recombinant ectodomain after proline-stabilization are instead due to the increased durability of the prefusion conformation of Env.

Leu568 and Thr569 are 99.8% and 98.9% conserved among all of the 6,599 Env sequences curated in the LANL HIV database (**S.Fig. 7**), likely reflecting their functional importance during the transition from prefusion-to-postfusion and their insulation from antibody-mediated selective pressure. This high degree of sequence conservation at positions 568 and 569 prompted us to evaluate whether our 2P stabilization strategy might also be effective outside the context of the CH848 10.17DT Env. A panel of multi-clade HIV-1 and SIV Envs (**S. Table 2**) was selected and the yield of recombinantly expressed prefusion trimer after transient transfection of a 2P-stabilized SOSIP construct was compared to that of a non-2P-stabilized SOSIP construct (**Fig. 5**). Overall, 2P stabilization improved the yield of soluble Env trimers from HIV-1 clades B and C, subgroup 02_AG and even the SIVcpz Env that was tested from CAM13K. However, the degree to which 2P stabilization enhanced Env yield was highly variable for the constructs that were evaluated. B41 and JRFL Envs exhibited fairly modest ∼2-fold increases in yield, whereas fold-increases for CAM13K, T250-4 and CH505 Envs could not be reliably calculated due to an inability to purify the non-2P-stabilized construct. These findings suggest that 2P stabilization is a broadly applicable approach to stabilizing the prefusion conformation of Env, albeit to varying degrees depending on the Env variant in question. Furthermore, the 2P mutations are compatible for addition to a variety of alternative Env stabilization strategies, including the SOSIP, DS and F14 mutations (**S. Table 2**).

## DISCUSSION

Conventional vaccination efforts have thus far been unable to elicit broadly neutralizing antibodies, which are correlated with protection against HIV-1 acquisition. These failures have prompted the development of new mosaic immunogens and sequential immunization strategies, the latter of which are intended to guide the immune system from germline targeting through the process of antibody maturation(34, 35). These proposed vaccination regimens require the production of numerous, diverse Env immunogens, many of which are recalcitrant to recombinant expression despite modification with previously reported stabilization strategies. Here we report a novel set of proline-stabilization mutations which facilitate the purification of antigenically diverse Envs by enhancing the durability of the prefusion conformation.

By targeting the region in gp41 that connects the central helix to the rest of heptad repeat 1 (HR1), the 2P mutations are predicted to disfavor the formation of the elongated alpha helices that drive the transition to the occluded-open and postfusion conformations. In this sense, they are mechanistically similar to the previously reported “HR1-redesigned” trimers(23), in which residues 548-568 of gp41 are replaced with a flexible linker. However, because the 2P mutations only require two amino acid substitutions at the highly conserved 568-569 positions, rather than removal of the HR1 and the introduction of an exogenous 8-10 amino acid loop, they present a less intensive engineering feat during the design and creation of new Env constructs. Moreover, we have shown that the 2P mutations can be used in conjunction with many existing stabilization strategies, including the SOSIP(18), DS(24) and F14(22) mutations. Additional studies are required to determine whether the 2P mutations are also compatible with alternative stabilization strategies, such as the recently described repair-and-stabilize approach(25, 36). Perhaps the most interesting opportunities to combine 2P with existing stabilization strategies will involve approaches that have already altered the residues at positions 568 or 569, such as the aforementioned HR1-redesigned trimers(23) or the MD2 trimer(37), which makes use of a L568D mutation to enhance expression of the BG505 Env.

Having developed this stabilization strategy and confirmed that it did not alter the antigenicity or structure of soluble Env trimers, we also show that its mechanism of increasing the yield of Env trimers is through enhancing the durability of the prefusion conformation, rather than increasing overall expression. During the relatively harsh, prolonged conditions of transient transfection, Env ectodomains that more reliably retain the prefusion conformation are presumably less likely to spontaneously and irreversibly trigger to the postfusion conformation and be lost as insoluble aggregates(30). In addition to facilitating the purification of recombinantly expressed prefusion Envs for use as biochemical reagents in the laboratory, the enhanced durability of 2P Envs may also have implications for future vaccine design and production efforts.

Many proposed vaccination regimens currently rely on extensive priming and boosting schedules, which would inevitably complicate vaccine adherence among the general population(38, 39). One reason for this approach is the difficulty associated with eliciting even autologous neutralizing antibodies(40, 41). By enhancing the durability of the prefusion conformation of Env, it may be possible to increase the residence time of Env immunogens in B cell germinal centers, theoretically mimicking the conditions of natural infection where natively expressed Env is continually present(15). Reliable presentation of the prefusion Env is thought to be strictly required in order to elicit a polyclonal, broadly neutralizing response since the UCAs of many bnAbs, such as DH270 and CH01, are only capable of recognizing Env in the prefusion conformation(28, 42). The enhanced durability observed in our thermostability and forced degradation assays suggest that 2P-stabilized Envs may not require the same restrictive cold-chain storage conditions that are necessary for unstabilized Env immunogens. In addition to the advantages that this poses in a laboratory environment, it is also a critical consideration when thinking about administering vaccines in regions that may lack sufficient infrastructure to reliably maintain a vaccine cold chain.

Finally, we also show that the 2P stabilization strategy is effective in multiple HIV-1 and SIV isolates. By introducing these mutations into a diverse panel of different Env ectodomains, we were able to enhance the yield of multiple prefusion trimers, albeit to differing degrees of success depending on the Env variant. This improvement in Env yield underscores the key importance of the HR1 region in trimer stability(23). But given the variant-dependent levels of efficacy that were observed with the 2P stabilization strategy, it is likely that other factors such as glycan networking or conformational heterogeneity within the gp120 subunit also have a significant impact on the overall yield of a given trimer(43). The complex chemical environment generated by the tertiary and quaternary structure of the Env trimer makes it difficult to pinpoint what precise features might be contributing to this variant-dependent efficacy. However, as mosaic and sequential vaccination strategies continue to be developed, it is our hope that having the 2P stabilization strategy as a broadly applicable means to enhance the yield of antigenically diverse prefusion trimers will prove useful for immunogen design.

## DATA AVAILABILITY

The cryo-EM map and corresponding atomic model have been deposited in the Electron Microscopy Data Bank (EMDB) and the Protein Data Bank (PDB) under accession codes EMD-28608 and PDB ID: 8EU8. All flow cytometry data are available upon request.

## AUTHOR CONTRIBUTIONS

D.W., Z.M., B.H.H., P.A., K.O.S. and B.F.H. designed research. D.W., Z.M., B.T., K.J., O.A., R.P. and M.B. performed research. D.W. and Z.M. analyzed data. D.W., K.O.S. and B.F.H. wrote the manuscript with input from all authors.

## ACKNOWLEDGMENTS

We would like to thank Aja Sanzone, Alecia Brown and Beth Bryan for their assistance with cell culture. We would like to thank Rory Henderson for helpful discussions regarding the putative mechanisms of 2P stabilization. We would also like to thank Mitchell Martin, Jessica Martinson and Bhavna Hora of the DHVI Sequencing Core. Flow cytometry was performed in the Duke Human Vaccine Institute Research Flow Cytometry Facility (Durham, NC). This work was supported by the Consortia for HIV/AIDS Vaccine Development grant UM1AI144371 from the Division of AIDS, NIAID as well as R01 AI050529 (B.H.H.), R37 AI 150590 (B.H.H) and NIAID grant # 5T32AI007392-32 (D.W.).

## MATERIALS AND METHODS

### Protein production and purification

Plasmids encoding for Env ectodomains (CH848 10.17DT DS-SOSIP, CH848 10.17DT DS-SOS-2P, CH848 10.17DT DS-SOSIP-2P, CAM13 Q171K DS-SOSIP, CAM13 Q171K DS-SOSIP-2P, B41 SOSIP, B41 SOSIP-2P, JRFL SOSIPv6, JRFL SOSIPv6-2P, T250-4 DS-SOSIP, T250-4 DS-SOSIP-2P, CH505w24 F14 DS-SOSIP and CH505w24 F14 DS-SOSIP-2P) were mixed with a plasmid encoding for furin at a ratio of 4:1. All Env constructs contained a C-terminal HRV3C cleavage site, a TwinStrepTag and an 8xHisTag. These plasmid mixtures were transfected into FreeStyle 293-F cells (Thermo Fisher) using polyethylenimine. Transfected supernatants were harvested and filtered five days post-transfection. Because CH848 Envs do not bind to PGT145, all CH848 constructs were purified by StrepTactin resin (IBA). All other Envs were purified by PGT145 affinity chromatography, as described previously(20). After affinity chromatography, Envs were further purified by size-exclusion chromatography using a Superose 6 Increase 10/300 GL column (Cytiva) in 2 mM Tris pH 8.0, 200 mM NaCl, 0.02% NaN_3_.

Plasmids encoding for the heavy and light chains of mAbs used for ELISA or flow cytometric analysis (CH65, N6, DH270 UCA, DH270.6, PGT151, PGT128, VRC26.25, 2F5, 17b, 19b, RM19R, DH1029, 2G12, F39F, PGT125, F105 and A32) were combined at a ratio of 1:1 and used to transiently transfect Expi293 cells (Thermo Fisher) with ExpiFectamine (Thermo Fisher). Transfected supernatants were harvested and filtered five days post-transfection and antibodies were purified using Protein A resin (Thermo Fisher). Eluted antibodies were then buffer exchanged into PBS.

### ELISA

Env protein containing a C-terminal StrepTag was bound in wells of a 384-well plate, which were previously coated with streptavidin (Thermo Fisher Scientific) at 2 μg/ml and blocked with PBS containing 4% (w/v) whey protein, 15% normal goat serum, 0.5% Tween-20, and 0.05% sodium azide. Proteins were incubated at room temperature for 1 hour, washed with PBS and 0.1% Tween-20, then mAbs were added in serial dilutions beginning at 25 μg/ml. Antibodies were incubated at room temperature for 1 hour, washed, and binding was detected with goat anti-human HRP (Jackson ImmunoResearch) and TMB substrate (Sera Care Life Sciences).

### Cryo-EM sample preparation and data collection

Purified CH848 10.17DT DS-SOSIP-2P was diluted to a final concentration of 1.8 mg/mL in 10 mM Tris pH 8.0. To prevent interaction of the trimer with the air-water interface during vitrification, the sample was incubated in 0.085 mM n-dodecyl β-D-maltoside (DDM). A 3.5 μL drop of protein was deposited on a Quantifoil-1.2/1.3 grid (Electron Microscopy Sciences) that had been glow discharged for 10 seconds using an easiGlow Glow Discharge Cleaning System (PELCO). After a 30 second incubation in >95% humidity, excess protein was blotted away for 2.5 seconds before being plunge frozen into liquid ethane using a Leica EM GP2 plunge freezer (Leica Microsystems). Frozen grids were imaged using a Titan Krios (Thermo Fisher) equipped with a K3 detector (Gatan). Data were collected using the Gatan Latitude software.

### Cryo-EM data analysis and refinement

Movies were imported into cryoSPARC v3.3.1 (44) and aligned using patch-based motion correction. Patch-based CTF estimation was then performed before 5,780,212 particles were selected using a non-templated blob picking strategy. Junk particles were removed by 2D classification, leaving a stack of 683,406 particles that were subjected to iterative rounds of ab initio volume calculation and heterogeneous 3D classification, leaving a final stack of 111,026 particles. These particles yielded a 3.95 Å reconstruction after performing asymmetrical (C1) non-uniform refinement, with the resulting map exhibiting C3 symmetry. Non-uniform refinement(45) was then performed again using the same particle stack and applying C3 symmetry which improved the resolution of the resulting reconstruction to 3.73 Å. This map was then subjected to post-processing using DeepEMhancer(46). A full description of the cryo-EM data processing workflow can be found in **S. Fig. 4**. Chain A, D, E, H, I and 1 from PDB ID: 6UM7 were used as a starting model that was docked into the sharpened map and re-modeled through iterative rounds of building and refinement in Coot(47), PHENIX(48) and ISOLDE(49).

### gp160 cell-surface expression and characterization

293-F cells (ThermoFisher, cat #R79007) were diluted to 1.25 × 10^6^ cells/mL and seeded to 12-well plates 2-3 hours before transfection. Transient transfection of DNA plasmids was performed with jetPRIME transfection reagent (Polyplus, cat #101000046) following the manufacturer’s instruction. Transfected cells were cultured in an incubator at 37 °C with 8% CO_2_ and shaking at 125 rpm for 24 hours before flow cytometry staining. 24 hours after transfection, 293-F cells were harvested, counted and then were rinsed with 1% BSA/PBS and pelleted at 500 *g* for 5 minutes. Next, cells were resuspended to a density of 1 × 10^6^ cells/mL in 1%BSA/PBS and 50,000 cells were aliquoted to each well of U-bottom 96-well plates. An equal volume of recombinant anti-HIV-1 Env antibodies at 4 μg/mL were added to cells to reach a working concentration of 2 μg/mL. Antibodies were incubated with cells at 4 °C for 30 minutes. Cells were then washed once with 150 μL 1% BSA/PBS and then incubated with 50 μL Goat anti-Human IgG Fc secondary antibody PE (ThermoFisher, cat #12-4998-82) at a final concentration of 2.5 μg/mL in 1% BSA/PBS. After a 30 minute incubation at 4 °C while protected from light, cells were washed once with 1x PBS and incubated with 100 μL LIVE/DEAD Fixable Aqua Dead Cell Stain (ThermoFisher, cat# L34966, 1:1000 in PBS) for 20 minutes at room temperature, protected from light. Next, cells were washed once with 100 μL 1% BSA/BSA, then resuspended in 50 μL 1% BSA 2mM EDTA 1% PFA in PBS. Flow cytometric data were acquired on an iQue 3 high-throughput flow cytometry system (Sartorius). Data were analyzed using FlowJo v10 (FlowJo).

### Freeze/thaw thermostability analysis

Purified CH848 10.17DT DS-SOSIP and CH848 10.17DT DS-SOSIP-2P were diluted to 0.2 mg/mL in 2 mM Tris pH 8.0, 200 mM NaCl, 0.02% NaN_3_. Samples were rapidly frozen in liquid nitrogen and thawed by incubation at 30 °C for 5 minutes. Thermostability measurements were collected using a Tycho NT.6 by increasing the temperature from 35 °C to 95 °C at a rate of 30.0 °C/minute. Tm was determined as the inflection temperature using the Tycho Nanotemper data processing software.

### Forced degradation assay

250 μg aliquots of purified CH848 10.17DT DS-SOSIP or CH848 10.17DT DS-SOSIP-2P were incubated at 42 °C for 0 hours, 48 hours or 96 hours. Aliquots were then run over a Superose 6 Increase 10/300 GL column (Cytiva Life Sciences) to evaluate Env integrity. Area under the curve was calculated using UNICORN 7.0 Evaluation software (Cytiva Life Sciences).

## FIGURE LEGENDS

**Supplementary Figure 1:**
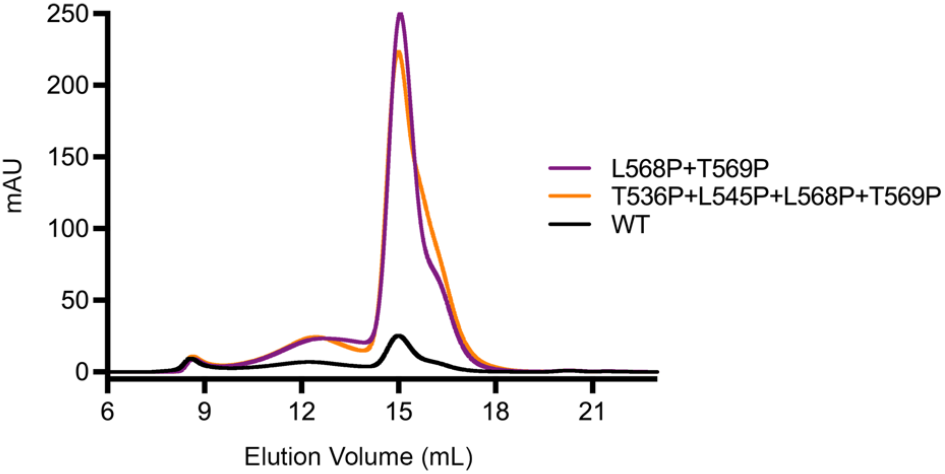
Evaluating effects of quadruple proline-substitution on Env homogeneity and yield. Size-exclusion chromatograms from a Superose 6 Increase column are shown. CH848 10.17DT DS-SOSIP is colored black (labeled “WT”), CH848 10.17DT DS-SOSIP L568P+T569P is colored purple and CH848 10.17DT DS-SOSIP T536P+L545P+L568P+T569P is colored orange. The curves from CH848 10.17DT DS-SOSIP and CH848 10.17DT DS-SOSIP L568P+T569P are the same as those shown in **Fig. 1C**.

**Supplementary Figure 2:**
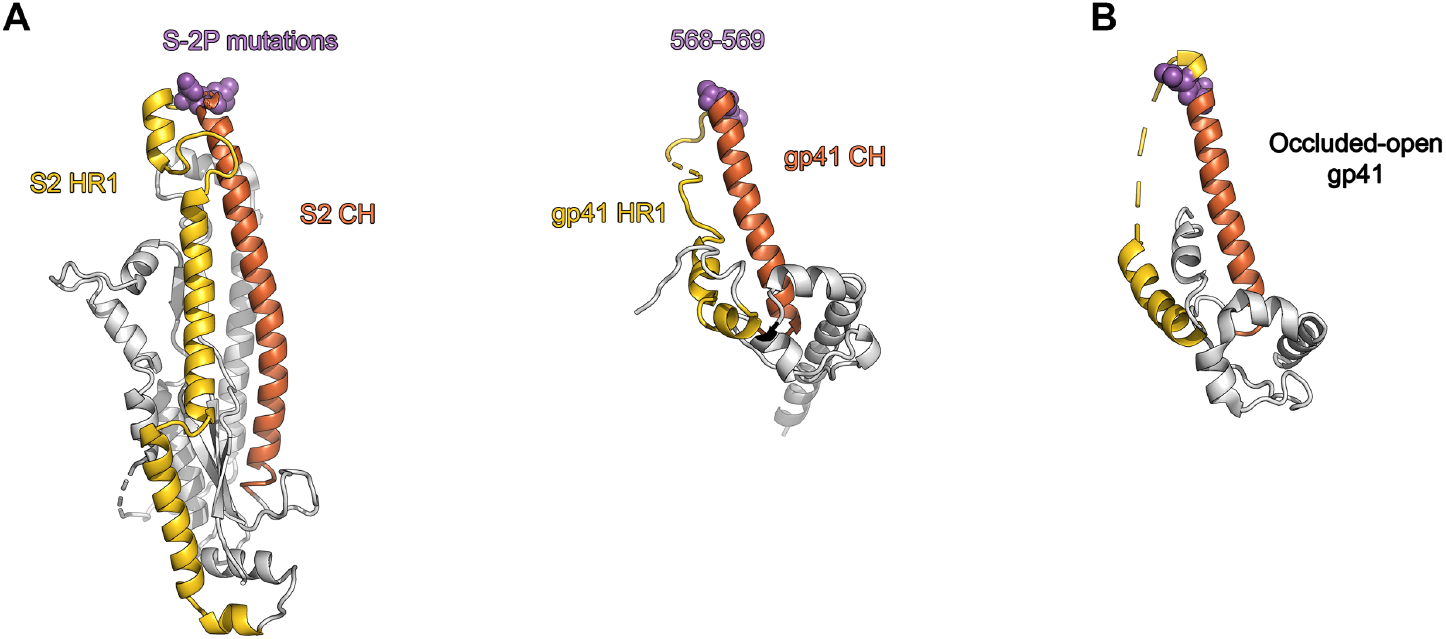
Structural comparison of CoV S-2P mutations and Env 2P mutations. (**A**, *left*) A monomer of the S2 subunit of the HCoV-HKU1 Spike in the prefusion conformation (PDB ID: 5I08) is shown as a ribbon diagram, with the heptad repeat 1 (HR1) colored yellow, the central helix (CH) colored orange, and the residues that are altered in the S-2P mutations shown as purple spheres. (**A**, *right*) A monomer of the gp41 subunit in the prefusion conformation (PDB ID: 6VZI) is shown as a ribbon diagram, with the HR1 colored yellow, the CH colored orange, and residues 568-569 shown as purple spheres. (**B**) A monomer of the gp41 subunit in the occluded-open state (PDB ID: 6CM3) is shown as a ribbon diagram, colored according to panel **A**.

**Supplementary Figure 3:**
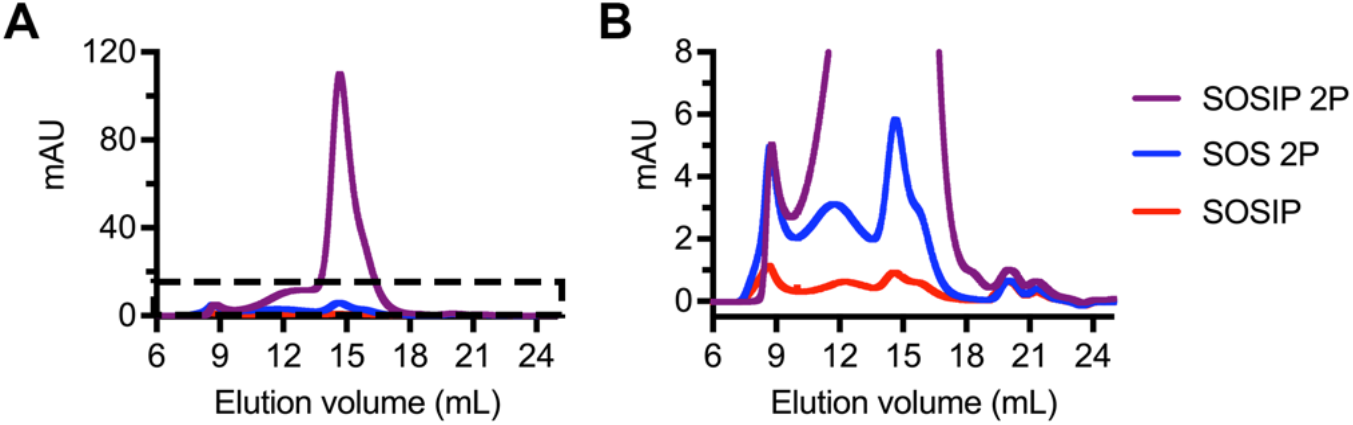
The I559P mutation and the 2P mutations act synergistically to boost CH848 10.17DT Env yield. (**A**) Size-exclusion chromatograms from a Superose 6 Increase column are shown. CH848 10.17DT DS-SOSIP is colored red (“SOSIP”), CH848 10.17DT DS A501C+T605C+2P is colored blue (“SOS 2P”) and CH848 10.17DT DS-SOSIP-2P is colored purple (“SOSIP 2P”). The dashed box denotes the zoomed-in view that is shown in panel **B**.

**Supplementary Figure 4:**
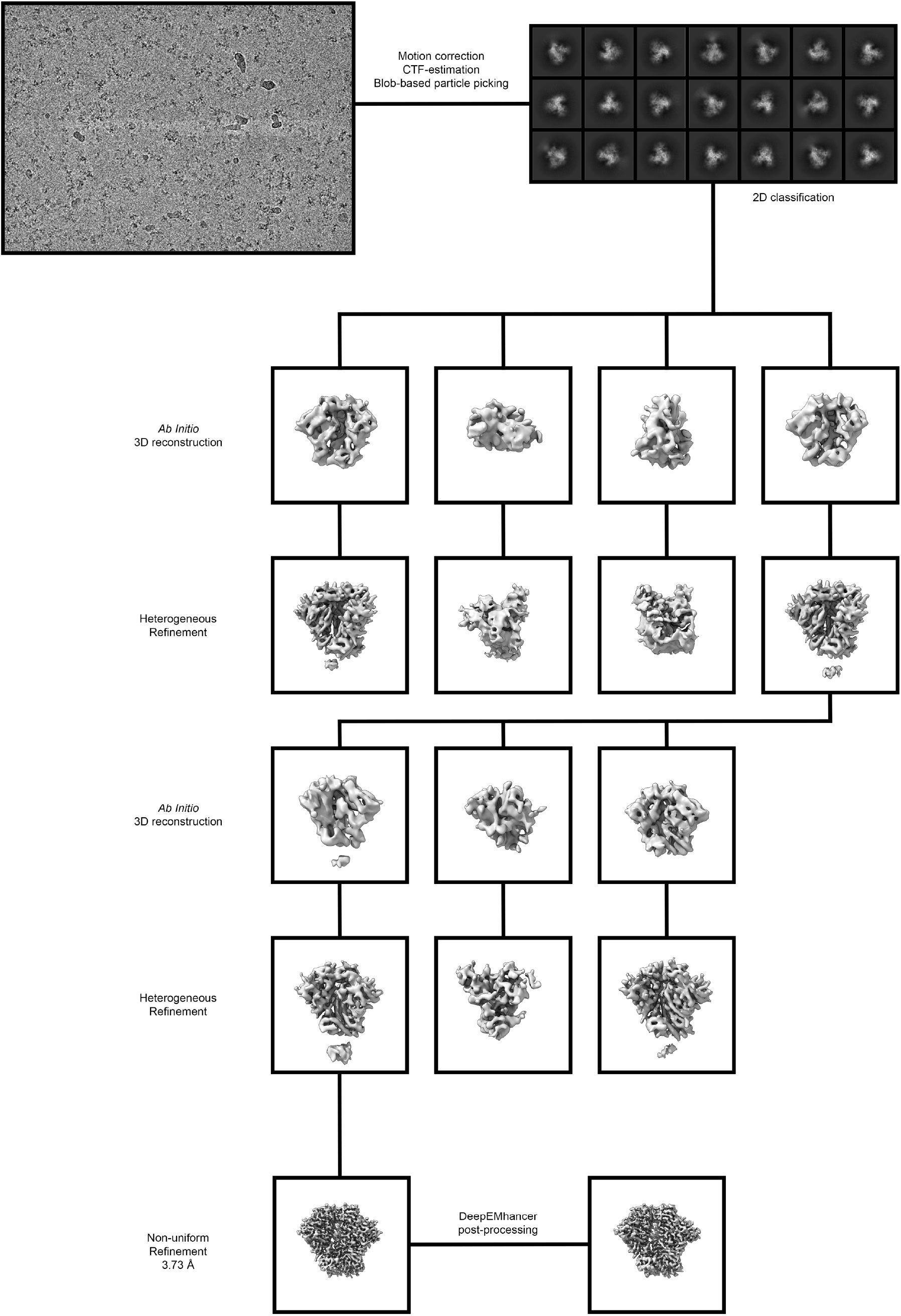
Cryo-EM data processing workflow.

**Supplementary Figure 5:**
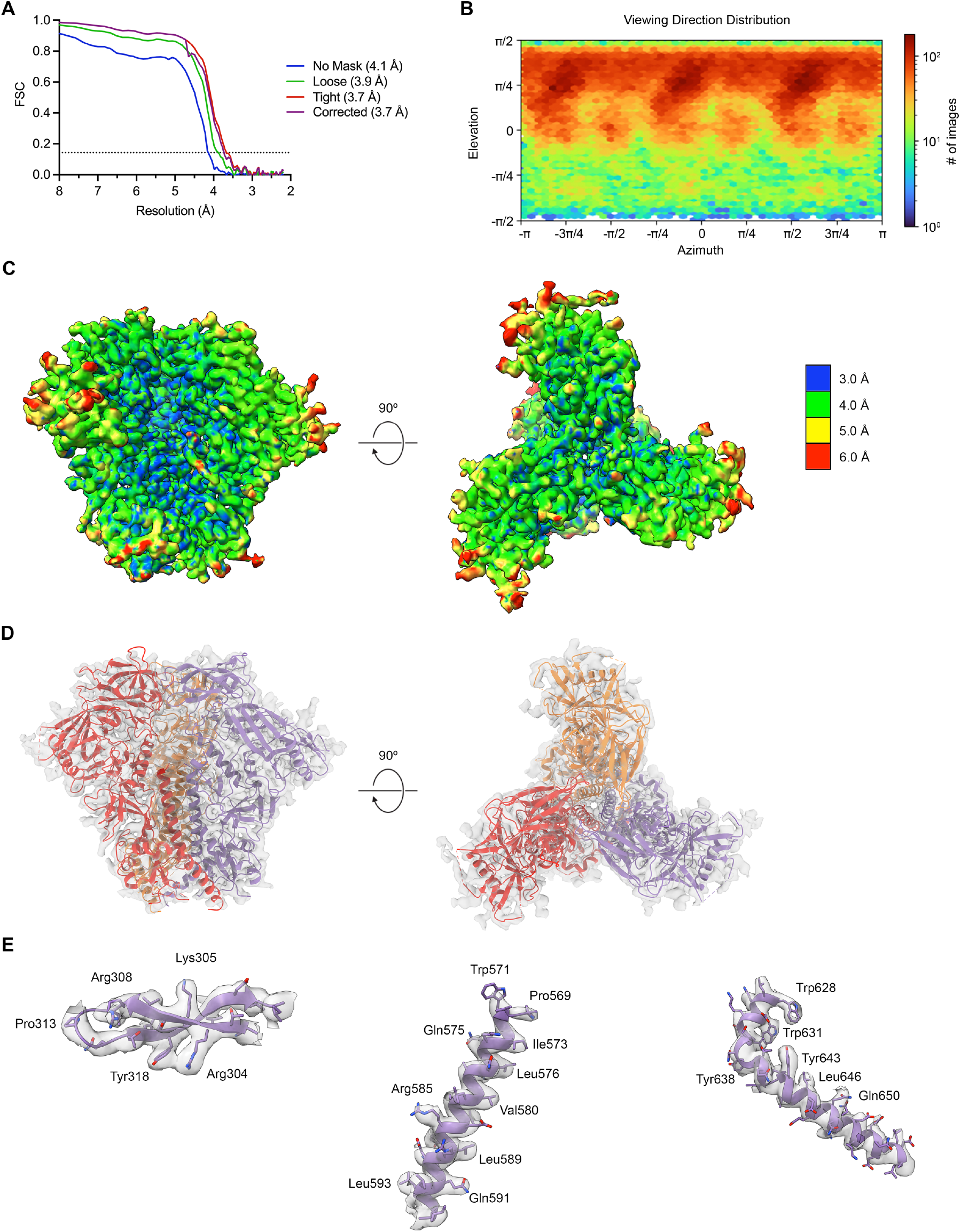
Cryo-EM validation. (**A**) FSC curves for different masking strategies employed by cryoSPARC v3 are plotted. The dotted line represents an FSC value of 0.143. (**B**) The viewing direction distribution plot, generated in cryoSPARC v3, is shown for the CH848 10.17DT DS-SOSIP-2P reconstruction. (**C**) Side (*left*) and top (*right*) views of the 3.73 Å CH848 10.17DT DS-SOSIP-2P reconstruction are shown, colored according to local resolution. (**D**) The cryo-EM map is shown as a transparent surface with the corresponding atomic model shown as a ribbon diagram, colored by protomer. (**E**) Portions of the cryo-EM map are shown as a transparent surface with the atomic model shown as a purple ribbon diagram. Side chains are shown as sticks with oxygen atoms colored red and nitrogen atoms colored blue.

**Supplementary Figure 6:**
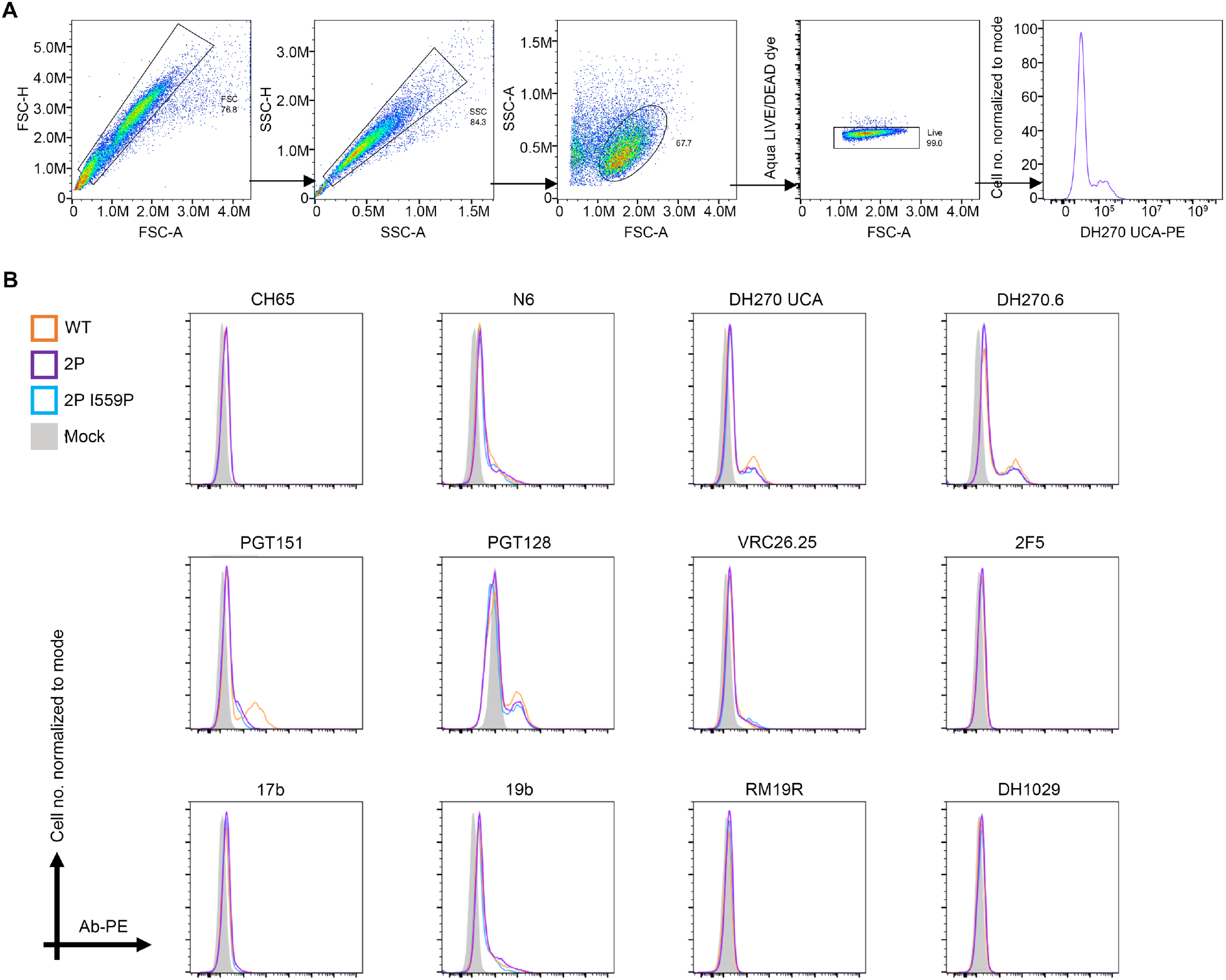
Flow cytometric analysis of proline-stabilized CH848 10.17DT gp160s. (**A**) Transiently transfected 293-F cells were gated to exclude doublets, debris and dead cells. mAb binding was detected using a PE-labeled goat anti-human IgG Fc secondary antibody. (**B**) Raw histograms for each mAb that is displayed in **Figure 4A** are shown. CH848 10.17DT gp160 curves are colored orange, CH848 10.17DT 2P gp160 curves are colored purple, CH848 10.17DT 2P I559P gp160 curves are colored blue and untransfected curves are colored gray. The number of cells, normalized to mode, is plotted on the y-axis and PE signal is plotted on the x-axis.

**Supplementary Figure 7:**
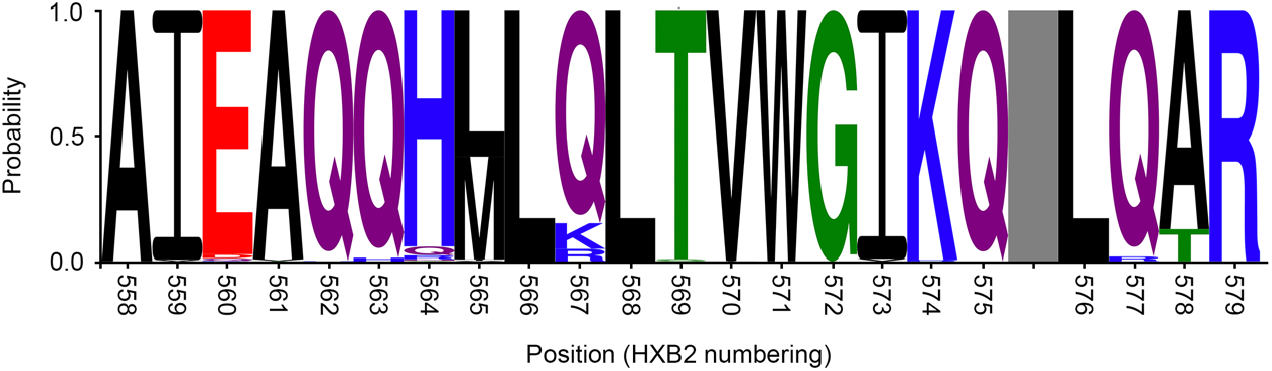
Conservation of residues Leu568 and Thr569. A WebLogo plot, calculated using 6,599 Env sequences curated in the LANL HIV database, is shown for positions 558-579. Residues have been colored according to their chemical properties (polar = green, neutral = purple, basic = blue, acidic = red, hydrophobic = black).

**Supplementary Table 1:**
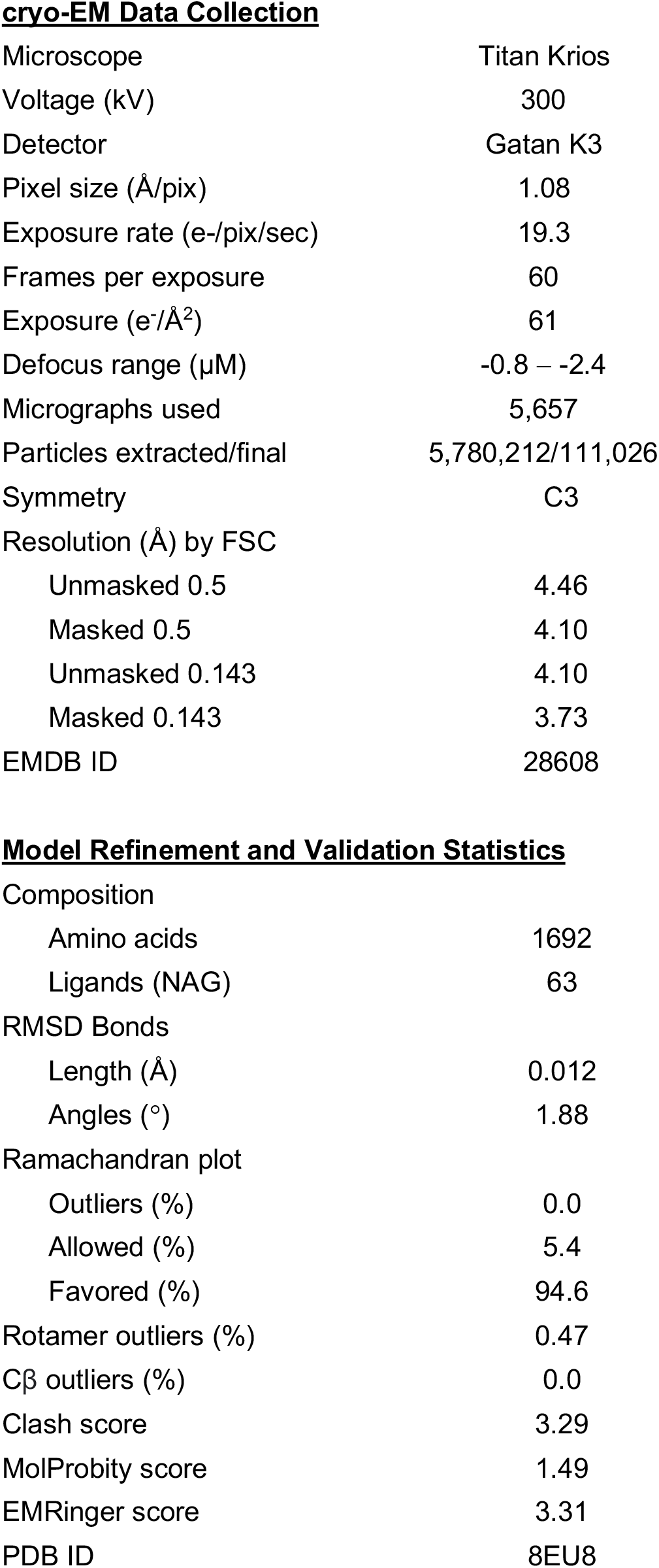
Cryo-EM collection and refinement statistics.

|  |  |
| --- | --- |
| Microscope | Titan Krios |
| Voltage (kV) | 300 |
| Detector | Gatan K3 |
| Pixel size (Å/pix) | 1.08 |
| Exposure rate (e<sup>-</sup>/pix/sec) | 19.3 |
| Frames per exposure | 60 |
| Exposure (e<sup>-</sup>/Å<sup>2</sup>) | 61 |
| Defocus range (μM) | -0.8 – -2.4 |
| Micrographs used | 5,657 |
| Particles extracted/final | 5,780,212/111,026 |
| Symmetry | C3 |
| Resolution (Å) by FSC |  |
| Unmasked 0.5 | 4.46 |
| Masked 0.5 | 4.10 |
| Unmasked 0.143 | 4.10 |
| Masked 0.143 | 3.73 |
| EMDB ID | 28608 |
| **Model Refinement and Validation Statistics** |  |
| Composition |  |
| Amino acids | 1692 |
| Ligands (NAG) | 63 |
| RMSD Bonds |  |
| Length (Å) | 0.012 |
| Angles (°) | 1.88 |
| Ramachandran plot |  |
| Outliers (%) | 0.0 |
| Allowed (%) | 5.4 |
| Favored (%) | 94.6 |
| Rotamer outliers (%) | 0.47 |
| Cβ outliers (%) | 0.0 |
| Clash score | 3.29 |
| MolProbity score | 1.49 |
| EMRinger score | 3.31 |
| PDB ID | 8EU8 |

**Supplementary Table 2:** Env panel characteristics.

| Abbreviated Name | Full Name | Clade/Subgroup | GenBank Accession Number | Additional mutations (excluding 2P) |
| --- | --- | --- | --- | --- |
| CH848 10.17DT | CH848.3.d0949.10.17 | C | KX217749 | SOSIP.664*, DS<sup>†</sup>, DT (N133D+N138T), chimeric BG505 gp41<sup>‡</sup> |
| CAM13K | SIVcpzCAM13K | SIVcpz | AY169968 | SOSIP.664, DS, Q171K |
| B41 | 9032-08.A1.4685 | B | EU576114.1 | SOSIP.664 |
| JRFL | JRFL | B | U63632.1 | SOSIPv6<sup>§</sup> |
| T250-4 | T250-4 | 02_AG | MW507842 | SOSIP.664, DS |
| CH505 | CH505w24 | C | n/a | SOSIP.664, DS, F14<sup>#</sup> |

\*SOSIP.664 = A501C+T605C+I559P, R6 optimization of furin cleavage site, truncation after residue 664
<sup>†</sup>DS = I201C+A433C
<sup>‡</sup>chimeric BG505 gp41 = Beginning at position 490, residues from BG505 Env were substituted for the native Env sequence
<sup>§</sup>SOSIPv6 = SOSIP.664 + E64K + A316W + A73C + A561C + E49C + L555C
<sup>#</sup>F14 = V68I+A204V+V208L+V255L

